# A novel emaravirus identified in maple with leaf mottle symptoms by deep sequencing

**DOI:** 10.1101/2020.10.05.325928

**Authors:** Artemis Rumbou, Thierry Candresse, Susanne von Bargen, Carmen Büttner

**Author notes:** Correspondence Artemis Rumbou.

## Abstract

The full-length genome of a novel *Emaravirus* has been identified and characterized from sycamore maple (*Acer pseudoplatanus*) - a tree species of significant importance in urban and forest areas - showing leaf mottle symptoms. RNA-Seq was performed using RNA preparations from a symptomatic and a symptomless maple tree. Purified double-stranded cDNA from each sample were used for RNA-Seq analysis on the Illumina HiSeq2500system and 14-198 MB data/sample of 100 bp-long paired-end sequence reads were generated. The sequence assembly and analysis revealed the presence of six RNA segments in the symptomatic sample (RNA1: 7,075 nt-long encoding the viral replicase; RNA2: 2,289 nt-long encoding the glycoprotein precursor; RNA3: 1,525 nt-long encoding the nucleocapsid protein; RNA4: 1,533 nt-long encoding the putative movement protein; RNA5: 1,825 nt-long encoding a hypothetical protein P5; RNA6: 1,179 nt-long encoding a hypothetical protein P6). Two independent HTS sequencing runs from the same symptomatic maple tree detected the same genome segments. For one of these sequencing runs the cDNA library was prepared using a primer targeting the conserved genome terminal region, known to be shared between emaraviruses genome segments and a high amount of sequence data was generated. We suggest, therefore, that the six identified genome segments represent the complete genome of a novel emaravirus from maple, which we tentatively name maple mottle-associated virus (MaMaV). RT-PCR assays were performed on symptomatic and non-symptomatic leaves of *A. pseudoplatanus* trees coming growing on two different locations in Berlin. MaMaV was only detected from symptomatic trees and all six RNAs were generally simultaneously detected. Non-symptomatic samples were consistently negative for MaMaV. These results suggest that MaMaV might be the symptom inducing virus in the sampled trees. In the present state of the art, this is the first time an Emaravirus is described from maple and is fully genetically characterized.

## Introduction

During the last decade numerous novel DNA and RNA viruses (Villamor et al., 2019) and viroids (Hadidi et al., 2020) have been uncovered in herbaceous as well as in woody hosts by applying a wide range of HTS methods for virus detection and discovery (Hadidi et al. 2016; Olmos et al., 2018; Villamor et al., 2019). While the focus of plant virology has mainly been narrowed to crop and fruit species, forest virology has also gained significant improvement due to the employment of HTS. Recently, a birch virome was unravelled revealing a complex of novel and known viruses (Rumbou et al. 2020), while a novel badnavirus associated with the birch leaf-roll disease was identified and characterized (Rumbou et al., 2018). European mountain ash ringspot-associated virus has been detected in new hosts like *Karpatiosorbus × hybrida* in Finland (von Bargen et al., 2020), *Sorbus intermedia* (von Bargen et al., 2019) and *Amelanchier* spp. (von Bargen et al., 2018). A novel emaravirus has been identified in mosaic-diseased Eurasian aspen (*Populus tremula*) (von Bargen et al., 2020). In earlier studies, a graft-transmissible agent causing leaf mottle in maples was reported, while rod-shaped particles approx. 300 nm-long and genomic material from tobamoviruses were observed by electron microscopy (EM) (Führling and Büttner, 1998). The present study aimed to determine through HTS viral agent(s) affecting maples with leaf symptoms in the urban forests in Berlin that have not been adequately characterized by conventional methods.

Viral diseases on maple species (*Acer sp*.) have been on the agenda of plant virologists for decades (Cooper, 1979). Atanasoff (1935) was probably the first to describe a “yellow mottle-mosaic” symptom in *A. negundo* and *A. pseudoplatanus* in Japan and Europe, possibly related with virus presence. Brieley (1944) observed chlorotic or ring mottle in *A. saccharum* in North America, and a similar symptom was reported by Ploaie and Macovei (1968) in Romania. Szirmai (1972) described a “mosaic mottling with chlorotic - tending to yellow ochre - spots” in *A. negundo* and *A. pseudoplatanus* in Hungary, while Cooper (1979) confirmed this symptomatology in maples in the UK. At the same time, mechanical transmission of the so-called “maple leaf perforation virus” from naturally infected maples to beans (*Phaseolus vulgaris*) was described by Subikova (1973), but without visualizing particles.

The graft-transmissibility of the agent causing leaf mottle in maples was reported (Führling and Büttner, 1998), while rod-shaped particles approx. 300 nm-long and genomic material from tobamoviruses were observed by electron microscopy (EM) in symptomatic leaves. Rod-shaped particles were also detected in young *A. saccharum* seedlings with chlorotic spots and mottle symptoms (Lana et al., 1980), which were attributed to tobacco mosaic virus (TMV) based on their serological and biological properties. Isometric particles of 26-30 nm diameter were detected in maple trees from Turkish urban areas (*A. negundo, A. pseudoplatanus, A. campestre*) exhibiting mottling, mosaic, leaf deformation and lateral shoot formation (Erdiller, 1986). These particles were attributed to arabis mosaic virus (ArMV), cucumber mosaic virus (CMV) and sowbean mosaic virus (SoMV). However, to date, *Acer sp*. has never been unambiguously described as a host for any well-characterized viral agent and data of the Virus-Host database also confirm that (Mihara et al., 2016).

Maples are abundantly found in European forests and urban parks, with the most common species *A. pseudoplatanus* (sycamore) and *A. platanoides* (Norway maple) representing a natural component of birch (*Betula* sp.) and fir (*Abies* sp.) forests (Gibbs and Chen, 2009). Several *Acer* species provide valuable timber and are the main sources of maple sugar and maple syrup (Bingelli, 1993). Damages on the trees have been regularly attributed to fungi with most harmful being *Verticillium* wilt, sooty bark disease caused by *Cryptostroma* species (Wulf and Kehr, 2009) as well as *Eutypella parasitica* causing trunk infections (Brglez et al., 2020). Phytoplasma-associated witches-broom disease has been reported in *A. negundo* (Kaminska and Suwa, 2006) as well as in japanese maple, *A. palmatum* (Li et al., 2012). Maple trees exhibiting virus-like symptoms were regularly observed in forests around Berlin as well as in other regions of Germany during the last 40 years (Bandte et al., 2008). Considering the ecological and economical importance of maples, we aimed to fill in the gap in maple’s pathology by uncovering viruses possibly affecting maple trees. As a result, the genome of a viral agent is fully characterized, while its association with the observed symptomatology is strongly suggested.

## 2 Materials & Methods

### 2.1. Plant Materials

In 2014, leaf samples exhibiting virus-like symptoms, including mottle and leaf deformation (Fig. 1), were collected from an *A. pseudoplatanus* tree in the Berlin-Grunewald urban forest [Acer+ (2014): symptomatic tree], where such symptoms have been monitored for at least two decades. Randomly selected leaf parts were pooled together and used for RNA extraction. A similar pool of leaves was obtained from a symptomless seedling [Acer- (2014): non-symptomatic tree]. In 2015, the same symptomatic tree was re-sampled and RNA was extracted from pooled leaves exhibiting symptoms [Acer+ (2015)].

**Figure 1.**
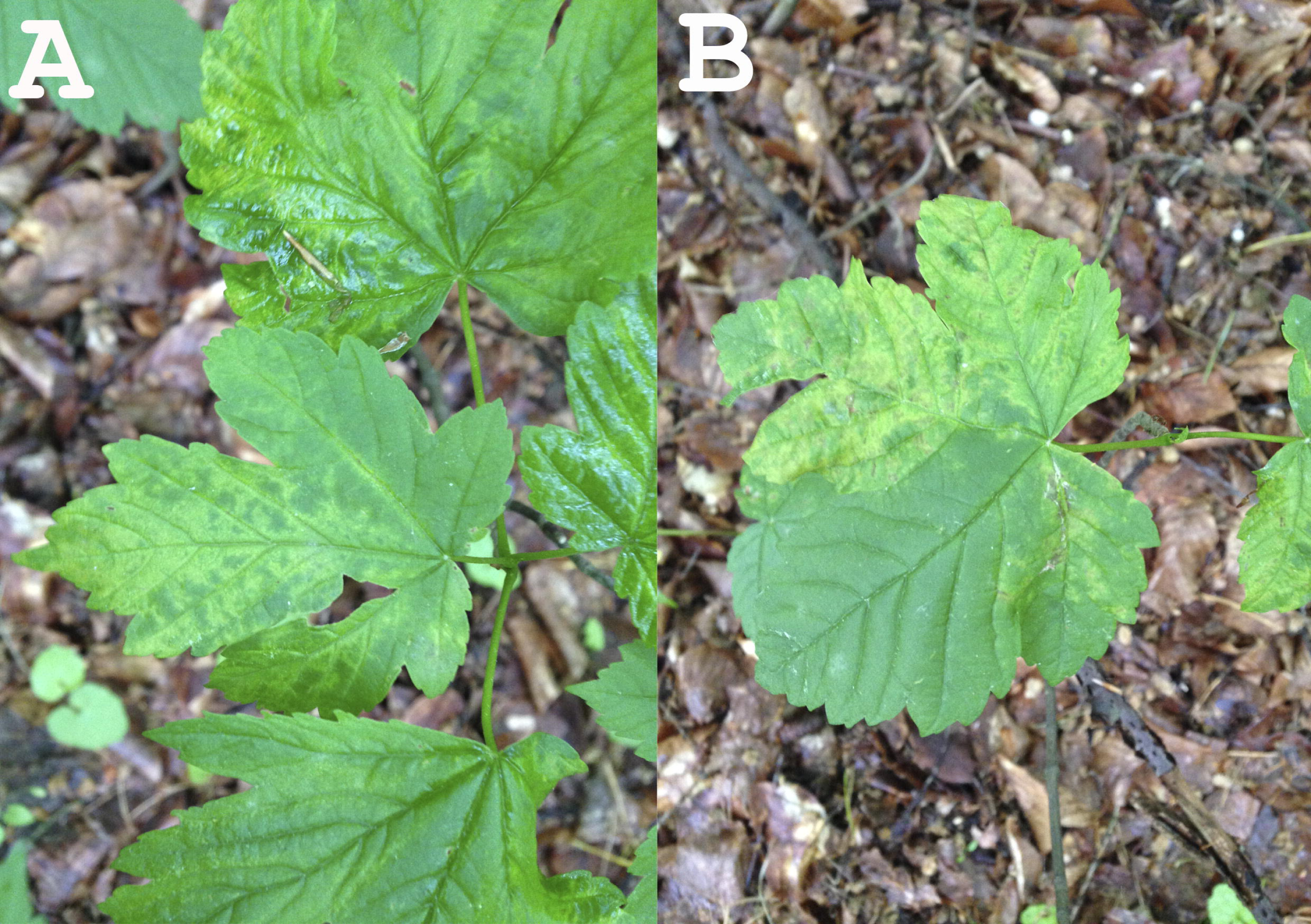
Leaf symptoms exhibited on leaves of the tested sycamore tree subjected to HTS (A: leaf mottle; B: leaf mottle and deformation).

For the investigation of the virus presence in the urban Berlin-Grunewald forest, leaves were collected from symptomatic and non-symptomatic maples in order to be used for RNA isolation and RT-PCR diagnostic assays. In total, 26 sycamore maple trees exhibiting symptoms similar to those of the Acer+ (2014) tree as well as 6 trees without symptoms were tested in 2015, 2016 or 2019 (Suppl. Table S1).

### 2.2. High-Throughput sequencing and sequence assembly

Total RNAs were isolated from 100 mg leaf tissue using the InviTrap® Spin Plant RNA Mini Kit (STRATEC Molecular, Germany), followed by removal of remaining DNA with rDNase according to the supplier protocol (Macherey-Nagel, Germany) and RNA purification using NucleoSpin® RNA Clean-up (Macherey-Nagel, Germany). Ribosomal RNA depletion was performed using the RiboMinus Plant Kit for RNA-Seq (Invitrogen). One to two micrograms of RiboMinus RNA of each sample were used for cDNA synthesis with the Maxima H Minus double-stranded cDNA synthesis Kit (Thermo Scientific) primed with random hexamers for samples Acer+ (2014) and Acer-(2014). For sample Acer+ (2015), the precipitated RNA was reverse transcribed into cDNA using the generic terminal primer PDAP213 (DiBello and Tzanetakis, 2013).

Two micrograms purified double-stranded cDNA from each sample were sent to BaseClear (Netherlands) for RNA-Seq analysis on the Illumina HiSeq2500system. FASTQ sequence files were generated using the Illumina Casava pipeline version 1.8.3. Initial quality assessment was based on data passing the Illumina Chastity filtering. Subsequently, reads containing adapters and/or PhiX control were removed using an in-house filtering protocol. The second quality assessment was based on the remaining reads using the FASTQC quality control tool version 0.10.0. HTS data processing and analysis were carried out either using CLC Genomics Workbench version 7.0.4. or the VirAnnot pipeline (Lefebvre et al., 2019). The quality of the FASTQ sequences was enhanced by trimming off low quality bases using the “Trim sequences” option of the CLC Genomics Workbench version 7.0.4. The quality-filtered reads were de novo assembled into contig sequences using CLC Genomics Workbench. Contigs annotation was carried out using BLASTn and BLASTx against the NCBI-GenBank databases. When needed, virus-related contigs were manually assembled into larger scaffolds and scaffolds polished by re-mapping of reads on the scaffolds using CLC Genomics Workbench.

### 2.3 Taxonomic analysis of the metagenome

The taxonomic content of the obtained datasets, as provided by the BLAST annotations, was visualized using MEGAN (Huson et al., 2016), in which the BLAST results were parsed to assign the best hits to appropriate taxa in the NCBI taxonomy. As a result, the taxonomical content (“species profile”) of the sample from which the reads were collected was estimated, with a particular focus on viral species (Fig 2; A– B).

**Figure 2.**
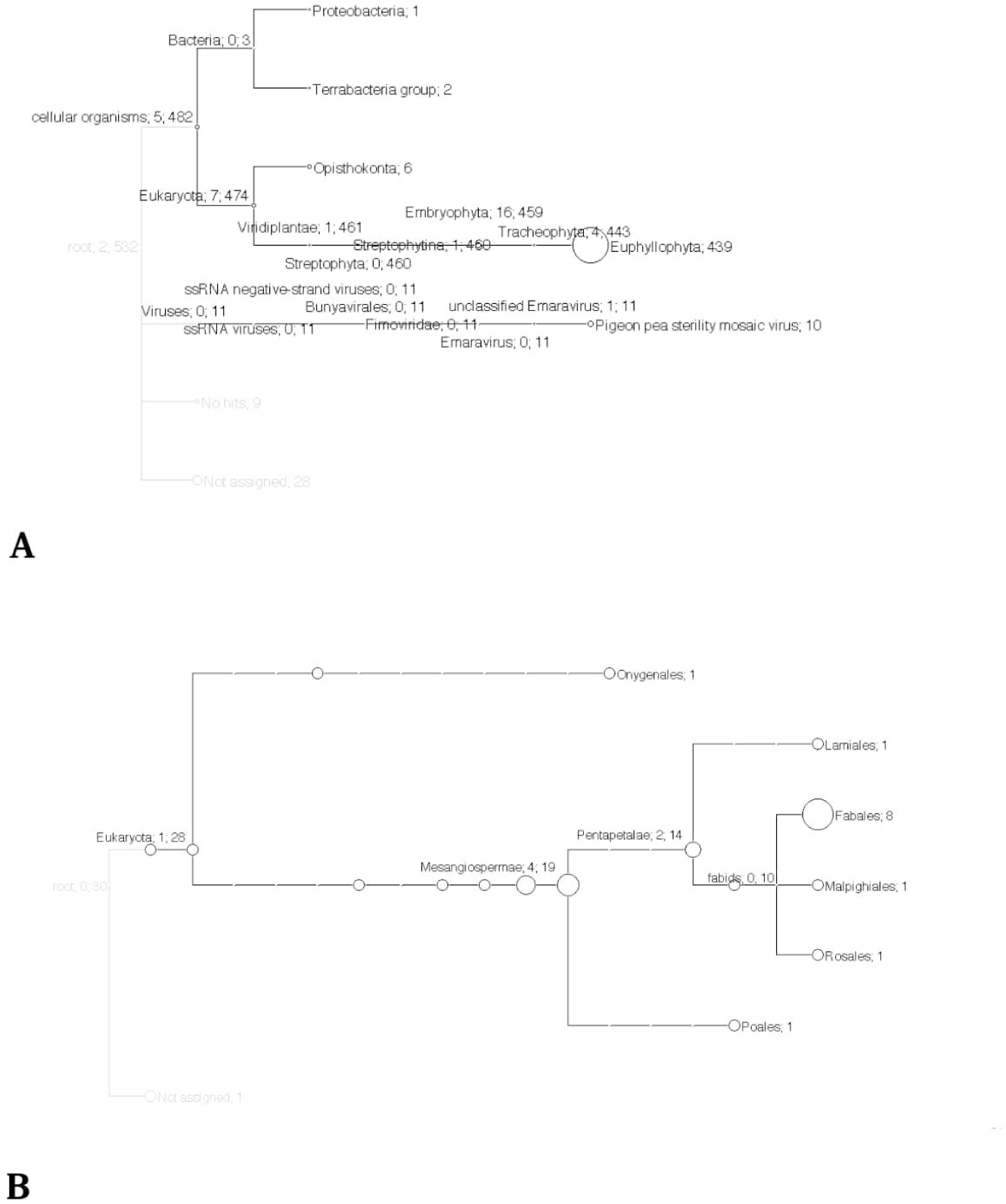
Taxonomical content of contigs assembled *de novo* from two maple RNA-Seq datasets (A) sample from symptomatic tree Acer+ (2014). (B) sample from non symptomatic tree Acer-(2014).

### 2.4 Validation of the presence of novel virus in maples

Total RNAs were isolated from 100 mg leaf tissue from 32 samples collected from symptomatic and non-symtomatic trees (Suppl. Table S1) using the extraction protocol described by Boom et al. (1990). Pooled samples of three to five leaves from different twigs of each tree were used. Symptomatic trees exhibited most often mottle, or mottle in combination with a mixture of other symptoms like flecking, chrolotic ringspots, vein banding, chlorotic line pattern, mosaic and/or leaf deformation.

The first-strand cDNAs were synthesized from 1 µg of total RNA in a 20 µl reaction volume of 1 x RT buffer (Thermo Scientific) containing 1 µM dNTPs mix, 100 U RevertAid Premium reverse transcriptase (Thermo Scientific), 20 U Ribolock RNase inhibitor (Thermo Scientific) and 100 pmol of random hexamer-oligonucleotides (biolegio). Subsequent PCR amplifications were conducted in a 50 µl volume of 1 x DreamTaq Buffer (Thermo Scientific) containing 0.2 µM dNTP mix, 0.25 U of DreamTaq DNA polymerase and 1 µM of each forward and reverse primer. The designed primer pairs targeted regions of each of the identified viral genome segments (Table 1). The thermal cycles were as follows: 2 min at 94 °C followed by 35 cycles at 94 °C for 30 s, 55 °C for 30 s, 72 °C for 30 s, with a final extension step of 72 °C for 5 min. The product lengths amplified for the different RNA segments are shown in Table 1.

**Table 1.**
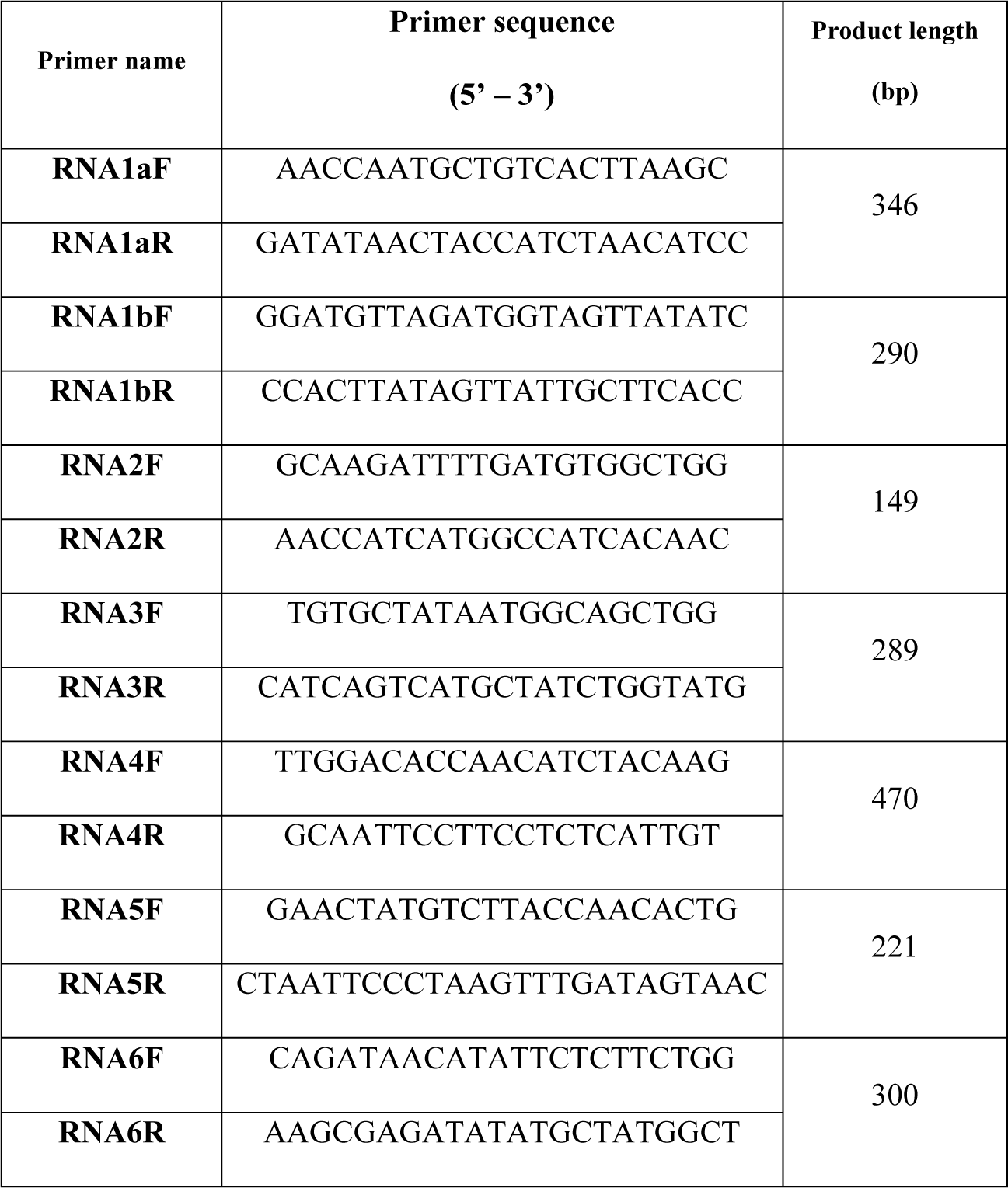
Features of primers used for the specific RT-PCR detection of the novel virus.

### 2.5 Sequence analysis and phylogenetic comparison of novel sequences

Multiple nucleotide or amino acid sequence alignments as well as pairwise sequence identity calculations were performed using AliView version 1.17.1 (Larsson, 2014). ORF finder at NCBI (https://www.ncbi.nlm.nih.gov/orffinder/) was used to identify open reading frames (ORFs) on assembled genome segments and identify the encoded proteins. All ORFs with 300 or more nucleotides were considered. For the phylogenetic comparisons of complete coding regions, the 21 established and tentative emaravirus species identified to date and represented in GenBank were used. Maximum likelihood (ML) trees were constructed with MEGA6 (Tamura et al., 2013) applying the Jones-Taylor-Thornton (JTT) substitution model for aminoacids. Robustness of nodes of the phylogenetic tree was assessed from 1,000 bootstrap replications, and values >70% were displayed for trees’ internal nodes.

To perform preliminary genetic divergence analysis, 40 RT-PCR products from 10 tested samples were directly submitted for Sanger sequencing (Macrogen) without previous cloning. They were amplified from three RNA segments; RNA1, RNA3 and RNA4. For RNA1, RT-PCR products were amplified in two different genome regions; the primer-pair RNA1aF/R generated PCR products of 311nt length (genome positions 3006 – 2756), while the primer pair RNA1bF/R generated 274 nt-long RT-PCR products located at nt positions 3317-3042. Evolutionary analyses were conducted in MEGA6 (Tamura et al., 2013). Genetic distance between sequences was assessed applying the Maximum Composite Likelihood model (Tamura et al., 2004), where the number of base substitutions per site between sequences was calculated.

## 3 Results

### 3.1 Quality analysis of FASTQ sequence reads and *de novo* analysis reveal a novel emaravirus from maple

RNA-Seq was performed in 2014 using RNA preparations from a symptomatic and a symptomless maple tree. 124 and 14 MB data/sample with approx. 35 Phred of average quality were generated for Acer+ (2014) and Acer-(2014), respectively (620,460 FASTQ reads for Acer+ (2014); 69,353 FASTQ reads for Acer-(2014)) (Table 2). For the sample Acer+ (2014) the *de novo* assembly of quality-filtered paired-end reads resulted in 532 assembled contigs. Analysis identified 14 contigs exhibiting significant aa identities to RNA sequences of several emaraviruses as assessed by BLASTp (7 contigs with identities to emaravirus-RNA1 assembling a 6,005 bp-long scaffold with missing genome parts within the sequence and at the ends; 3 contigs with identities to emaravirus-RNA2 assembling a 1,761 bp-long scaffold; one contig 1,085 bp-long with identities to emaravirus-RNA3; one 1,256 bp-long contig with identities to emaravirus-RNA4; one 1,277 bp-long contig with identities to emaravirus-RNA5; one 928 bp-long contig with identities to emaravirus-RNA6). All assembled contigs/scaffolds were missing sequences at the 3’ and 5’ ends. In the negative control sample Acer- (2014) none of the 30 contigs assembled showed any significant identities to emaraviruses or any other plant viruses. No other contigs from the Acer+ (2014) sample were identified as viral besides the 14 ones showing affinities with emaraviruses.

**Table 2.** Quality statistics of FASTQ sequence reads and *de novo* statistics for the three maple samples.

| Plant | FastQ<br>sequence reads | Sample Yield<br>(in MB) | *De novo* assembly |  |  |  |  |
| --- | --- | --- | --- | --- | --- | --- | --- |
|  |  |  | Total reads | Matched<br>reads | Total contigs | Average<br>contig<br>length(bp) | Emara-specific<br>contigs |
| **Acer+ (2014)** | 620,460 | 124 | 1,116,554 | 924,998 | 532 | 677 | 14 |
| **Acer- (2014)** | 69,353 | 14 | 123,424 | 115,795 | 30 | 984 | 0 |
| **Acer+ (2015)** | 850,283 | 198 | 1,700,566 | 1,496,745 | 2,206 | 646 | 8 |

To confirm the RNA-Seq results from the Acer+ (2014) sample and to complete the missing parts of the detected genome segments, the whole deep sequencing process was repeated in 2015 with leaves from the same tree. A ds-cDNA library was generated using the emara-specific terminal primer PDAP213, aiming to detect the missing genome ends. The RNA-Seq analysis provided a higher amount of sequence data (198MB), delivering 850,283 FASTQ sequence reads (av. quality score: ∼35 Phred) (Table 2). The *de novo* assembly resulted in a total of 2,206 contigs with an average length of 646 bp. BLASTp analysis identified 8 long contigs exhibiting significant aa identities to emaraviruses (3 with identities to emaraviruses RNA1, assembling a 6,907 bp-long scaffold missing the 5’ end; one 2,289 bp-long contig with identities to emaraviruses RNA2; one 965 bp-long contig with identities to emaraviruses RNA3; one short 235 bp-long contig with identities to emaraviruses RNA4, one 1,851 bp-long contig with identities to emaraviruses RNA5 and one 1,100 bp-long contig with identities to emaraviruses RNA6). The new contigs showed 100% nucleotide identity with the ones generated from the same tree in 2014. As such, they were assembled together with the 2014 contigs, providing the full-length sequence of the various RNA segments. Finally, a polishing step was performed to ensure genome completion and the absence of errors and to confirm genome ends. As a result, complete genomes for the six RNA segments were obtained (RNA1: 7,075 nt; RNA2: 2,289 nt; RNA3: 1,525 nt; RNA4: 1,533 nt, RNA5: 1,825 nt; RNA6: 1,179 nt). The full-length genomic sequences of the maple emaravirus RNA segments have been deposited in NCBI under accession numbers MT879190 -MT879195.

### 3.2 Taxonomic analysis of the metagenome

The contigs resulting from the *de novo* assembly of the reads for each sample were used for the MEGAN analysis. For the symptomatic sample Acer+ (2014), out of the 532 contigs assembled, 474 belong to Eucaryota, most of them to clade *Euphylophyta* (Phylum: Streptophyta; Kingdom: Viridiplantae) — where *Acer sp*. is classified — and three belong to Bacteria (Fig 2A). The 11 viral contigs identified by MEGAN are attributed to the *Fimoviridae* family and ten are attributed to pigeon pea sterility mosaic virus (*Emaravirus, Fimoviridae*). In the case of the non-symptomatic sample Acer- (2014), none of the 30 assembled contigs is attributed to a viral species.

The taxonomic analysis performed by MEGAN show a high degree of consistency with the results delivered by BLASTp annotation and suggest the precence of an emaravirus in the tested samples. MEGAN results obtained regarding the taxonomic content of contigs assembled from the RNA-Seq reads are shown in Fig 2 (A-B) together with the number of contigs assigned to each taxon.

### 3.3 Genome structure and encoded proteins

Applying ORF Finder, 6 open reading frames (ORFs) were identified, one for each RNA segment. BLASTn and BLASTp annotation of the assembled scaffolds from the symptomatic maple tree revealed high BLAST scores with RNA segments from viruses of the genus *Emaravirus* (*Fimoviridae, Bunyavirales*).

ORF1 predicted on RNA1 is 2,305 aa-long and the encoded protein shows in BLASTp analysis significant aa identity with the viral replicase of 21 known emaraviruses (ORF1: positions 6966 -49 nucleotides, nt) ranging from 32 to 75% (Table 3). The putative RNA polymerase exhibits highest aa identity with that of rose rosette emaravirus (Accession number: QHZ99251.1; 74,4% aa identity).

**Table 3.** Pairwise comparison of sequence identities on the amino acid level of putative proteins (RdRp, GPP, N, MP, P5, P6) encoded by RNA1-RNA6 of the novel emaravirus from *Acer pseudoplatanus* with the homolog proteins of the established and putative members of the genus Emaravirus.

|  | **RdRp** | **GPP** | **N** | **MP** | **P5 55 kDa** | **P6 26 kDa** |
| --- | --- | --- | --- | --- | --- | --- |
| **Rose rosette emaravirus** | **74.40%**<br>QHZ99251.1 | **56.65%**<br>QID76023.1 | 51.77%<br>QIB97971.1 | 64.84%<br>QIB98058.1 | **44.81%**<br>QIB98219.1 | 42.04%<br>QJR96770.1 |
| **Aspen mosaic-associated virus** | 70.23%<br>CAA0079389.1 | 53.11%<br>CAA0079597.1 | 51.04%<br>CAA0079646.1 | 57.73%<br>CAA0079685.1 | - | 40.39%<br>CAA0079719.1 |
| **Actinidia emaravirus 2** | 69.66%<br>QEE82886.1 | 47.78%<br>QEE82887.1 | **55.63%**<br>QEE82888.1 | 63.31%<br>QEE82889.1 | 43.41%<br>QEE82890.1 | **44.16%**<br>QEE82891.1 |
| **Pistacia emaravirus** | 69.16%<br>QAR18002.1 | 52.07%<br>QAR18003.1 | 51.60%<br>QAR18004.1 | 62.33%<br>QAR18005.1 | 40.46%<br>QAR18006.1 | 32.43%<br>QAR18008.1 |
| **Fig mosaic emaravirus** | 68.67%<br>QBH72675.1 | 52.15%<br>QBK46595.1 | 54.04%<br>BAM13809.1 | **65.19%**<br>BAM13817.1 | 33.98%<br>YP_009237273.1 | 41.71%<br>BAM13854.1 |
| **Pigeonpea sterility mosaic emaravirus 1** | 53.58%<br>ANQ90714.1 | 46.56%<br>QBA83603.1 | 40.29%<br>CDX09880.1 | 62.64%<br>ANQ90777.1 | 42.92%<br>CUR49054.1 | 37.70%<br>ANQ90719.1 |
| **Pigeonpea sterility mosaic emaravirus 2** | 68.14%<br>QBA83607.1 | 51.32%<br>YP_009268865.1 | 53.29%<br>ALU34071.1 | 64.00%<br>ANQ90759.1 | 43.89%<br>QBA83611.1 | 37.70%<br>ANQ90763.1 |
| **Blackberry leaf mottle-associated virus** | 66.09%<br>AQX45473.1 | 52.22%<br>AQX45474.1 | 50.17%<br>AQX45475.1 | 56.32%<br>QBM15152.1 | - | 35.57%<br>AQX45477.1 |
| **European mountain ash ringspot-associated** | 48.70%<br>VFU05375.1 | 40.67%<br>YP_003104765.1 | 36.69%<br>SPN63240.1 | 35.24%<br>VFU05382.1 | - | - |
| **Actinidia chlorotic ringspot-associated virus** | 47.82%<br>YP_009507925.1 | 41.53%<br>YP_009507926.1 | 38.43%<br>YP_009507928.1 | 33.70%<br>YP_009507927.1 | - | - |
| **Lilac chlorotic ringspot-associated virus** | 48.92%<br>QIN85945.1 | 42.61%<br>QIN85946.1 | 40.69%<br>QIN85947.1 | 36.56%<br>QIN85948.1 | - | - |
| **Redbud yellow ringspot-associated emaravirus** | 47.27%<br>YP_009508083.1 | 40.96%<br>YP_009508087.1 | 37.91%<br>YP_009508085.1 | 35.85%<br>YP_009508084.1 | - | - |
| **Ti ringspot-associated emaravirus** | 37.26%<br>QAB47307.1 | 26.55%<br>QAB47308.1 | 30.82%<br>QAB47309.1 | 23.10%<br>QAB47310.1 | - | - |
| **Raspberry leaf blotch emaravirus** | 36.87%<br>YP_009237274.1 | 26.98%<br>YP_009237265.1 | 28.57%<br>YP_009237266.1 | 24.08%<br>YP_009237267.1 | 31.91%<br>YP_009237268.1 | - |
| **Palo verde broom virus** | 35.24%<br>AWH90165.1 | 26.42%<br>AWH90170.1 | 24.42%<br>AWH90176.1 | 27.00%<br>AWH90182.1 | - | - |
| **Jujube yellow mottle-associated virus** | 37.60%<br>QDM38999.1 | 28.39%<br>QDM39000.1 | 28.23%<br>QDM39001.1 | 24.53%<br>QDM39002.1 | - | - |
| **High Plains wheat mosaic emaravirus** | 34.97%<br>YP_009237277.1 | 28.17%<br>QGT41075.1 | 26.55%<br>YP_009237257.1 | 24.14%<br>QGT41077.1 | 33.33%<br>QGT41070.1 | - |
| **Camellia japonica-associated emaravirus 1** | 32.61%<br>QGX73503.1 | 26.60%<br>QGX73504.1 | 29.73%<br>QGX73505.1 | 19.93%<br>QGX73506.1 | - | - |
| **Camellia japonica-associated emaravirus 2** | 32.89%<br>QGX73507.1 | 25.93%<br>QGX73508.1 | 25.85%<br>QGX73509.1 | 21.07%<br>QGX73510.1 | - | - |
| **Perilla mosaic virus** | 32.07%<br>BBM96177.1 | 23.75%<br>BBM96178.1 | 32.58%<br>BBM96180.1 | 21.36%<br>BBM96181.1 | - | - |
| Common oak ringspot-associated emaravirus | 35.73%<br>LR828198 | 28.04%<br>LR828199 | 27.27%<br>LR828200 | - | - | - |
| Pea-associated emaravirus | - | - | 50.52%<br>QJX15716.1 | - | - | - |

ORF2 encoded on RNA2 is predicted to encode a protein of 646 aa (nt 1996-56). In BLASTp comparisons the encoded protein shows 26-57% aa identity with 21 other emaraviruses (Table 3). The highest aa identity is with the glycoprotein precursor (GPP) of rose rosette emaravirus (Accession number: QID76023.1; aa identity: 56,7%).

ORF3 encoded on RNA3 is predicted to encode a protein of 300 aa (nt 104-1006) with highest BLASTp aa identity with the nucleocapsid proteins of the 22 known emaraviruses (24-56%) (Table 3). The N protein is the only protein identified from pea associated virus, with which ORF3 shows relatively high identity (50,5 %). The ORF3-encoded protein shows highest aa identity with the N protein of actinidia emaravirus 2 (Accession number: QEE82888.1; aa identity: 55,6%).

ORF4 encoded on RNA4 is predicted to encode a protein of 363 aa (nt 1179-88) which shows very variable BLASTp aa identity levels with the movement proteins identified from 20 emaraviruses (19-65%) (Table 3). It exhibits highest aa sequence identity with the movement protein of fig mosaic virus (Accession number: BAM13817.1; aa identity: 65,2%).

The fifth predicted ORF is encoded on RNA5 and is predicted to encode a protein of 373 aa (nt 1526-93). The hypothetical protein encoded by RNA5 shows sequence homology (supported by E-values < than 0.01) with the corresponding protein detected from eight emaraviruses (30-43% aa identity) (Table 3). This putative protein shares highest identity with the P5 protein of rose rosette emaravirus (Accession number: QIB98219. 1; aa identity: 44,8%), a protein of unknown function. The rest of the emaraviruses do not have a homologous protein or such has not been identified yet.

Finally, the sixth ORF, encoded on RNA6 (nt 766-71), produces a 231 aa-long protein with 35-45% aa identity with that of other emaraviruses (Table 3). The highest aa identity is with the P6 protein from actinidia emaravirus 2 (Accession number: QEE82891.1; aa identity: 44,2%). Again, only eight from the already known emaraviruses are found to have a homologous RNA segment to the one identified here.

We suggest that the segments RNA1-RNA6 represent the complete genome of the novel virus (Figure 4). It is known that Emaraviruses may contain up to eight different genome segments (Elbeaino et al., 2018). The type species, *European mountain ash ringspot-associated emaravirus* from the host *Sorbus intermedia* was recently shown to have 6 and not 4 segments as previously reported (von Bargen et al., 2019). More lately, a novel emaravirus in Shiso named *Perilla mosaic virus* (PerMV) was found to consist of 10 RNA segments, each encoding a single protein in the negative-sense orientation (Kubota et al., 2020). In the case of the maple virus, however, two independent HTS samples from the same maple tree detected the same number of genome segments. Additionally, for the Acer+ (2015) sample, the cDNA library was prepared using a primer targeting the conserved genome terminal region, known to be shared among emaraviruses genome segments and a high amount of sequence data was generated. A screen for further genome segments containing these terminal regions was not successful and we therefore consider that it is unlikely to have missed additional RNA segments of the viral genome.

### 3.4 Phylogenetic analysis of the novel emaravirus

Phylogenetic relationships between the maple virus and the sequences of 22 emaraviruses known to date were estimated, based on amino acid sequences comparisons. Irrespective of the RNA segment investigated, the maple viral agent clusters consistently in subgroup a (according to von Bargen et al., 2020), together with -among others - rose rosette emaravirus, fig mosaic virus, the novel aspen mosaic-associated virus and the two pigeonpea sterility mosaic emaraviruses and actinidia emaravirus 2. The phylogenetic analysis performed here confirms the overall structure within the genus *Emaravirus* described by von Bargen et al. (2020) and enriches the tree with seven novel emaraviruses. Fig 3 (A-D) shows representative ML trees obtained using the putative polymerase, the glycoprotein precursor, the nucleocapsid and the movement proteins encoded by the genome segments RNA1, RNA2, RNA3 and RNA4, respectively. The maple emaravirus proteins P5 and P6 are homologous to those of eight different emaraviruses (Suppl. Fig 3, E-F). All predicted proteins cluster together with the respective ones from rose rosette emaravirus. The only exception is the N protein, which clusters still with the subgroup a, but with no closest relation to any of its members.

**Figure 3.**
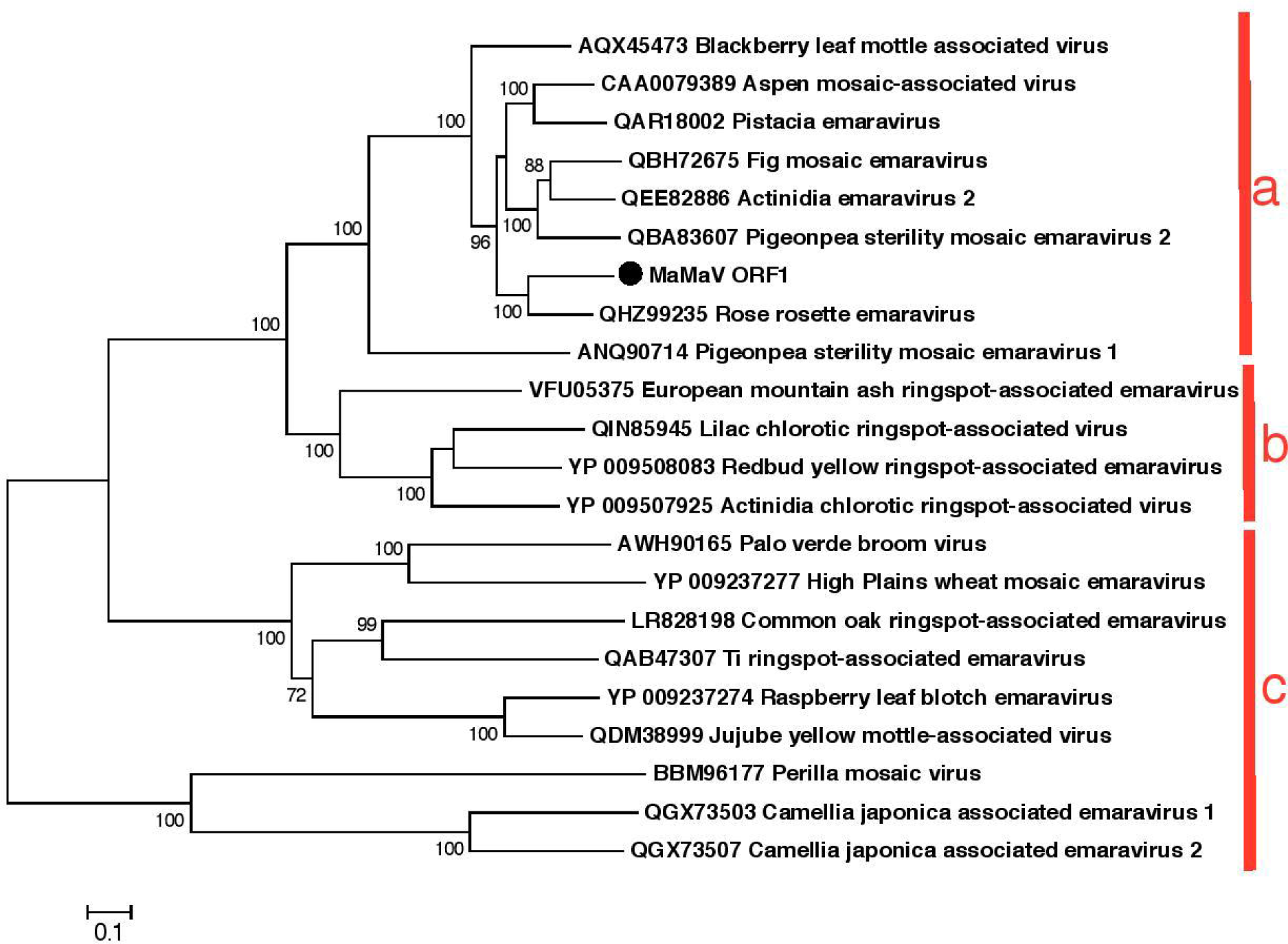
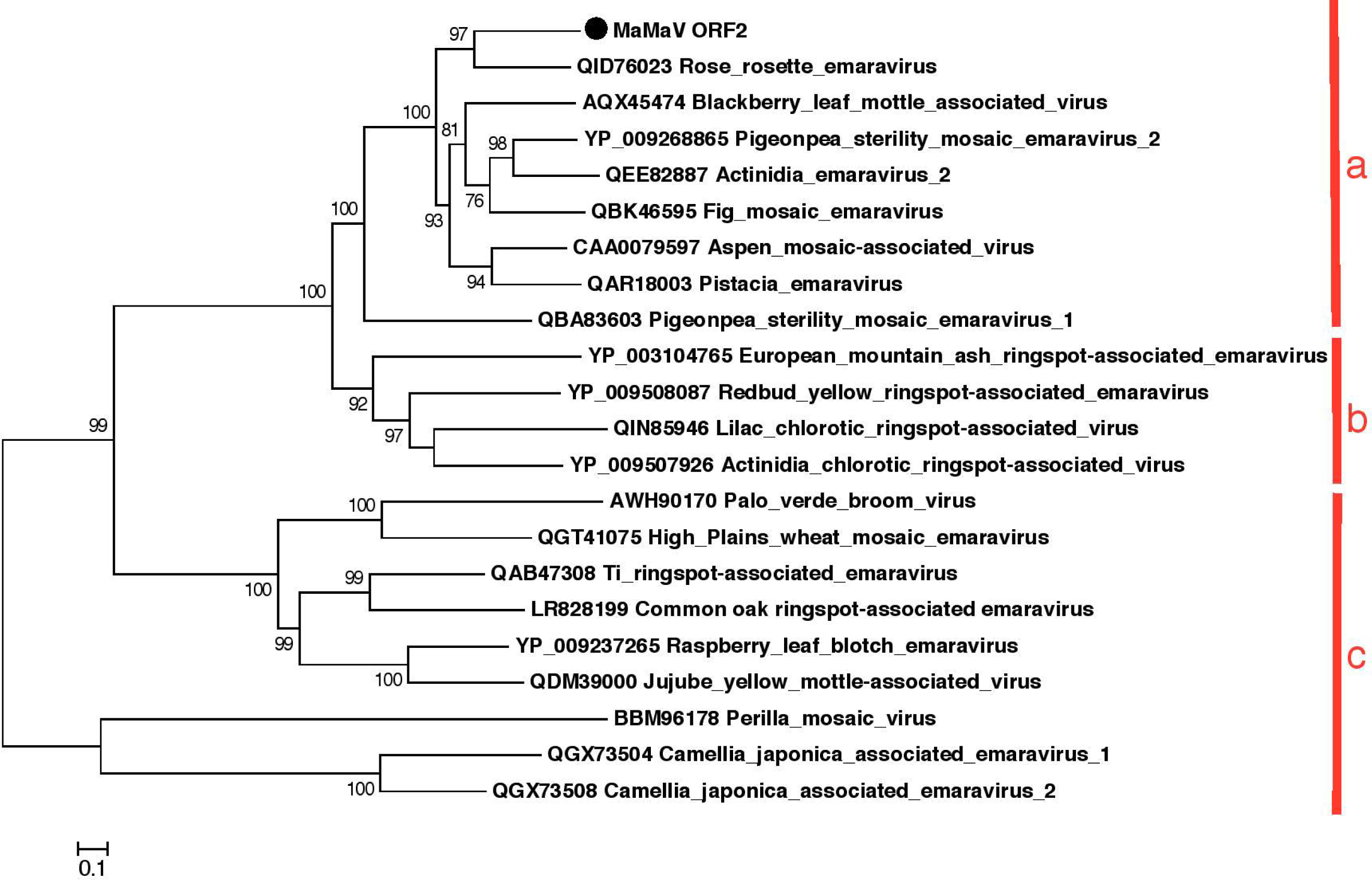
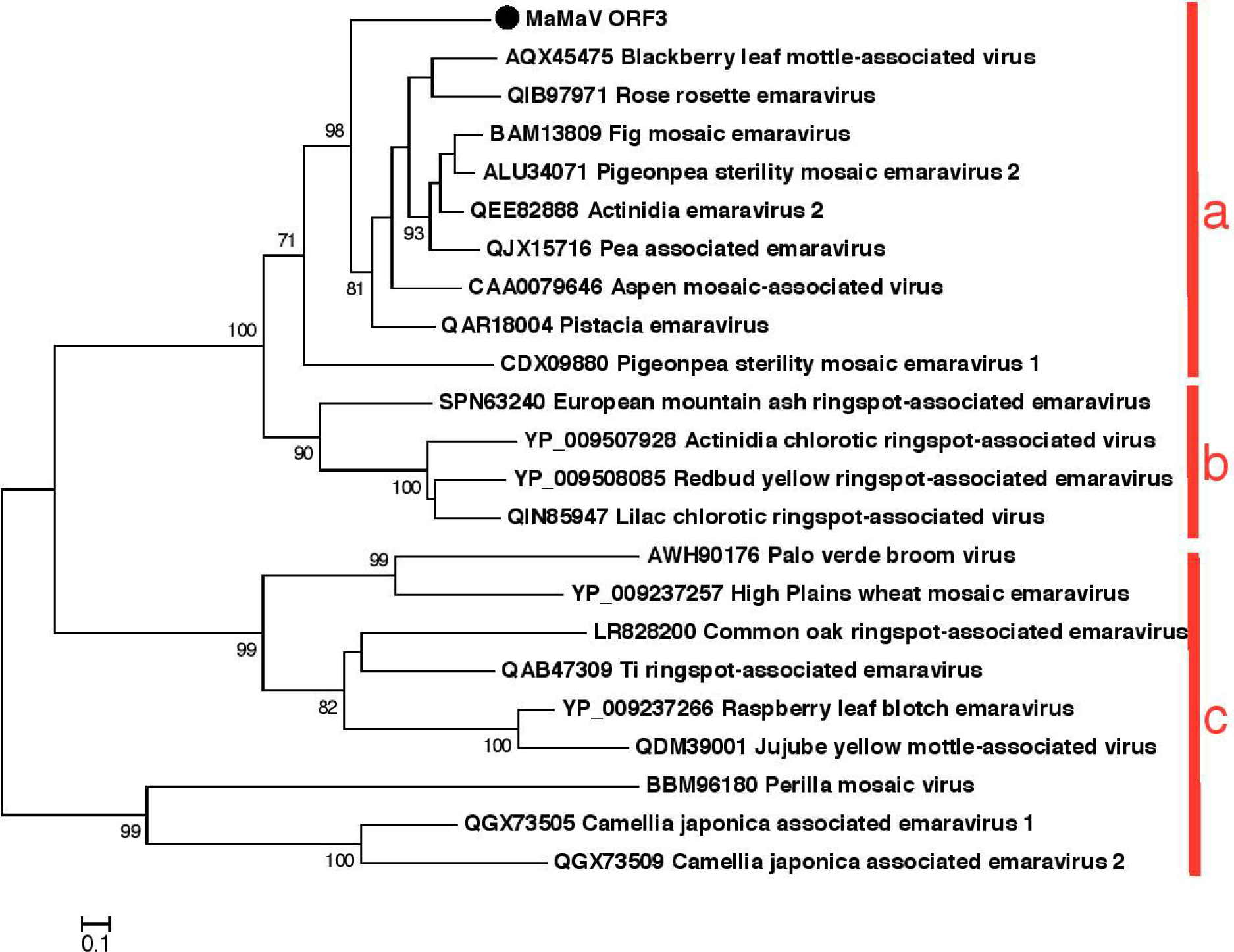
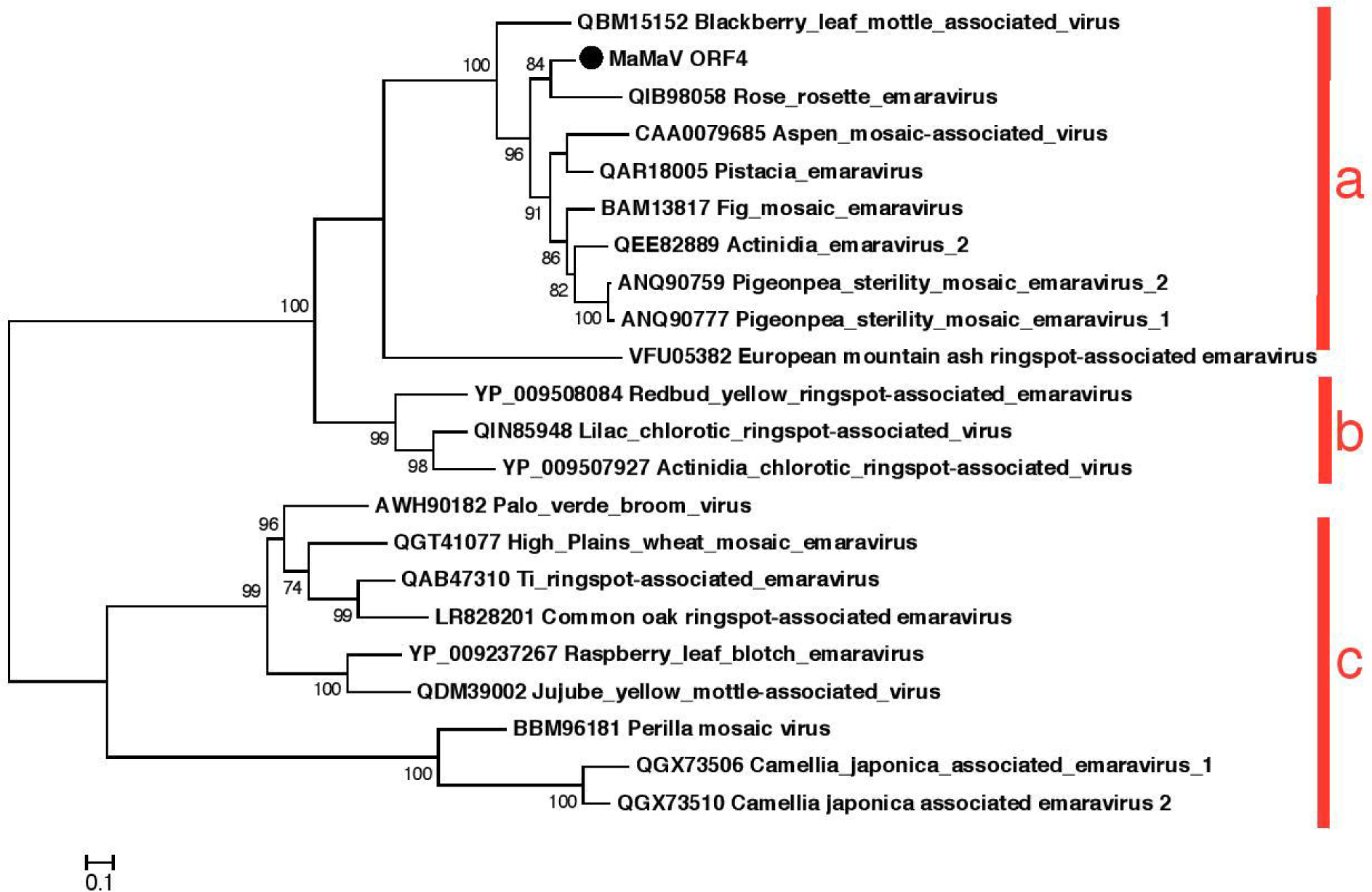
Phylogenetic trees reconstructed using the amino acid sequences of the RdRp (A), GPP (B), N (C) and MP (D) of the novel emaravirus. The tree was reconstructed using the Maximum Likelihood method and the statistical significance of branches was evaluated by bootstrap analysis (1,000 replicates). Only bootstrap values above 70% are indicated. The scale bar represents 10% amino acid divergence. Subgroups a, b and c within the genus *Emaravirus* described in Elbeaino et al. (2018) are indicated at the right side of the trees.

**Fig 4.**
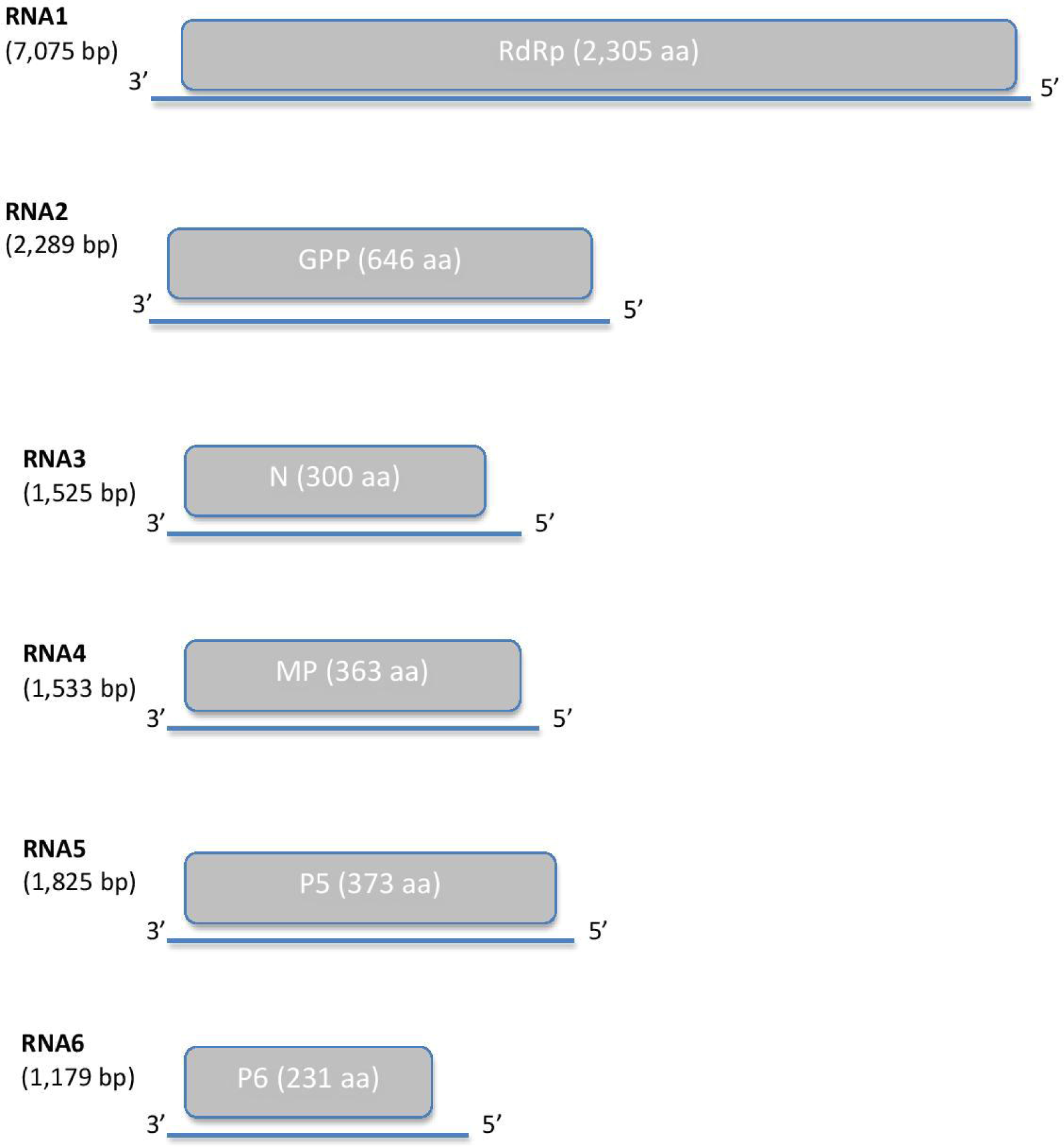
Schematic representation of the genome organization of the novel maple mottle-associated virus (MaMaV).

While the predicted proteins of the maple emaravirus exhibit significant aa homologies with the ones from other emaraviruses, they are clearly only distantly related with them. Taken together, these results demonstrate that the virus identified represents a new species in the genus *Emaravirus* and it is, therefore, tentatively named maple mottle-associated virus (MaMaV).

### 3.5 Validation of the RNA-Seq results and association of virus presence with symptom appearance

To confirm the presence of the identified segments in various maple samples, specific RT-PCR assays were performed using segment-specific primer pairs (Table 1). RT-PCR assays were performed on symptomatic and non-symptomatic *A. pseudoplatanus* trees coming from two different locations in Berlin-Grunewald and from grafted scions of that origin propagated in an experimental garden since 2017. The results are shown in Supplementary Table S1. MaMaV was only detected in symptomatic trees and all 6 RNAs were generally simultaneously detected. For a few samples, one of the six RNA segments failed to be detected (in sample E54934, segment RNA2; in samples E54936 and E54942, segment RNA1 tested with primer-pair RNA1bF/R - but both samples were tested positive with primer-pair RNA1aF/R - ; in sample E54946, segment RNA5). Because these cases were rare and all other RNA segments in those samples were successfully amplified, we suggest the negative results were due to the low concentration of respective RNA segments beyond detection limit of RT-PCR assay. Non-symptomatic samples were consistently negative for all MaMaV segments. These results suggest that the 26 positive-tested symptomatic maples were MaMaV-infected and that MaMaV could be the virus responsible for the identified virus-like symptoms.

It should be however underlined, that the tested trees showed a variability of symptoms, where mottle was mainly observed, but other -weaker or stronger-symptoms were also exhibited. Although mottle was the main and common symptom exhibited in the majority of infected trees, occasionally other virus-like symptoms, like chlorotic ringspots, line pattern or flecking and in some rare cases also leaf deformations were observed. More concretely, apart from mottle, in 2015 also flecking was recorded, while in 2016 chlorotic ringspots and line pattern were observed, although in both years samples were collected at the same time (1st of July; see Supplementary Table S1).

### 3.6 Preliminary results on genetic variability of three RNA segments

The evolutionary divergence between variants from different trees was estimated for three RNA segments. In this RNA segment low genetic diversity was revealed, with maxima of respectively 2% divergence for RNA1aF/R RT-PCR products and 1,1% for RNA1bF/R RT-PCR products. Sequences were identical in two and in 13 of the pairwise comparisons for the RNA1aF/R and RNA1bF/R RT-PCR products, respectively. However, all sequences differed from the corresponding variant generated from the original sequence from Acer + (2014) by 0.3-1.6% and by 0.4-1.1% in the cases of the RT-PCR products amplified by primer-pair RNA1aF/R and RNA1bF/R, respectively.

Sequences of RT-PCR products for the RNA3 segment were 274 nt-long and placed at nt position 734-468. In the RNA4 segment RT-PCR products of 431 nt-length were amplified and sequenced (nt position 835-405 in RNA4). No genetic diversity was shown between those sequences and all were identical with the corresponding segment from the original sample Acer+ (2014).

Based on these results, evidence is provided that the diversity in the local Berlin-Grunewald MaMaV population is generally low and that different level of genetic diversity may characterize its different genome segments.

## Discussion

The 6 newly identified RNA segments are attributed to a novel *Emaravirus* species based on the following: (a) The multipartite genome is composed of six single-stranded RNA molecules; (b) All six RNAs share a fully conserved stretch of 13 nucleotides at their 5′ and 3′ termini; (c) Each segment of the genome encodes a single protein, which shows sequence identity with homologous proteins of other emaraviruses; (d) In all phylogenetic trees generated with amino acid sequences, MaMaV is only distantly related phylogenetically to the emaraviruses currently represented in the GenBank fulfilling the current species demarcation criteria of emaraviruses of more than 25 % aa divergence of RNA1-RNA3 encoded proteins (Elbeaino et al., 2018). To our knowledge, for first time an emaravirus is described from maple and is fully genetically characterized.

The HTS method applied not only led to the discovery and genetic characterization of the novel emaravirus; it unraveled at the same time the maple virome, in terms of identifying the exhaustive collection of nucleic acid sequences deriving from viral agents. The maple virome of the concrete symptomatic tree tested is found to be very simple, as it includes a single variant from a single virus. The lack of virome complexity is rather surprising, when we consider obtained HTS results from other wild as well as cultivated woody hosts. Complex virome was revealed in birch, where up to five virus variants were identified in the transcriptome of individual trees (Rumbou et al., 2020). Concretely in birch, it has been demonstrated that not only multiple viral species but also diverse variants of the same virus may accumulate in single trees (Rumbou et al., 2016; Rumbou et al., 2020). Similarly, in single peaches multi-viral infection of up to six viruses and viroids was detected (Jo et al., 2018). The multiplicity of viral infections was also shown in metagenome samples from single plum trees (Jo et al., 2020). A possible explanation for the single viral infection in the maple tree could be the age of the tree; it was very young (approx. three years-old) when sampled, thus it was shortly exposed to viral pathogens.

The novel virus was only present in the tested symptomatic maple trees in Berlin, while it was not detected in non-symptomatic trees. This suggests that MaMaV presence is associated with the leaf symptoms identified in maples. Although the main observed symptom was mottle, other -weaker or stronger-symptoms were also exhibited. Whether MaMaV is the only causal agent for the different kind of symptoms or whether other viruses are also involved in the symptomatology needs to be further investigated. Furthermore, further efforts are needed to satisfy Koch’s postulates and firmly establish its causal role.

Virus-like symptoms apart from mottle have been also observed and reported earlier not only in Germany (Bandte et al., 2008; Büttner et al., 2013), but also in Romania (Ploiaie and Macovei, 1968), Hungary (Szizai, 1972), Turkey (Erdiller, 1986), North America (Cooper, 1979). Symptom descriptions such as “mosaic”, “chlorotic mottle”, “yellow-mottle”, “ring-mottle”, “mosaic mottling with chlorotic spots” were used by earlier researchers. Whether the newly identified virus might be associated with the symptoms described in these earlier studies remains to be determined, given in particular that electron-dense structures known as double-membrane-bound bodies (DMBs) with diameters differing within the range of 80 to 200 nm and typical of emaraviruses (Mielke-Ehret & Mühlbach, 2012) have never been reported in earlier electron microscopy samples from symptomatic maples, while more typical plant virus particles sometimes have. Furthermore, infection of maples by tobamoviruses reported by Führling and Büttner (1998) could not be confirmed by the present study. In that case, we suggest either that the samples observed by EM were contaminated during sample processing or that indeed tobamoviruses may also affect maples. It remains to be confirmed by future HTS analyses whether other viral agents may be present in symptomatic maples.

From the discovery of the first viral agent in maple thanks to the development of deep sequencing tools to its full biological characterization there is a long way to go (Massart et al., 2017). To investigate its potential for emergence, possible vectors and mode(s) of dispersal need to be determined. Several members of the genus *Emaravirus* are known to be transmitted by gall mites (Mielke-Ehret and Mühlbach, 2012; Hassan et al., 2017; Elbeaino 2018). In the present study, in several of the sampled trees damages from gall mites (*Aceria macrophylla, Eriophyes psilomerus*) as well as from leafhoppers were found. These arthropods can be considered as putative vectors, but the hypothesis that they may be involved in the emaravirus transmission needs to be studied. Regarding MaMaV’s impact on infected trees, we assume that the mottle symptoms exhibited in infected leaves may lead to reduced photosynthetic capacity and, consequently, to trees’ health deterioration. Based on the available diagnostic RT-PCR assays designed in the frame of the present study, the investigation of the agents’ distribution and its impact on the trees health can be estimated.

Until recently, only a low number of viral agents in forest trees had been characterized, due to two main reasons; (a) biased sampling based on a narrow focus on agricultural crops and fruit trees viruses (Büttner et al., 2013) and (b) a general bias against the identification of the most divergent genomes (Zhang et al., 2018). Emaraviruses are negative-sense single-stranded viruses that were until recently overlooked, not only when applying conventional methods, but also by metagenomic studies (Bejerman et al., 2020). The application of HTS resulted in the discovery of 13 emaraviruses during the last five years (Bejerman et al., 2020). Employing deep sequencing methodologies significant presence of emaraviruses in forests has been revealed. The last three years four emaraviruses were discovered in six forest or urban tree species; aspen mosaic-associated virus (AsMaV) in *Populus tremula* (von Bargen et al., 2020a), European mountain ash ringspot-associated virus (EMARaV) in *Sorbus intermedia* (von Bargen et al., 2019), *Karpatiosorbus × hybrid* (von Bargen et al., 2020b), and *Amelanchier* sp. (von Bargen et al., 2018), common oak ringspot-associated virus (CORaV) in *Quercus robur* (Rehanek et al. unpublished), and maple mottle-associated virus (MaMaV) in *Acer pseudoplatanus*. Based on existing knowledge on viral disease of fruit trees we suggest that viral agents might be a considerable stress factor for forests trees and may as well predispose affected trees to other more harmful stress factors (Büttner et al., 2013; 2015). Additional efforts on the field of forest virology are needed to provide concrete data on the magnitude of the tree damage due to viral infections.

## Data availability

The datasets generated for this study can be found in the NCBI database GenBank, accession numbers from MT879190 -MT879195.

## Authors Contributions

Conceptualization: Artemis Rumbou (AR), Carmen Büttner (CB). Data curation: AR, Thierry Candresse (TC). Formal analysis: AR, TC. Funding acquisition: CB. Investigation: AR, TC, Susanne von Bargen (SvB). Methodology: AR. Project administration: CB. Software: AR, TC. Validation: AR, TC, SvB. Visualization: AR, SvB. Writing – original draft: AR. Writing – review & editing: TC, CB, SvB.

## Acknowledgements

This work is partly accomplished in the frame of the COST Action FA1407-DIVAS (Deep Investigation of Virus Associated Sequences)—‘Application of next-generation sequencing for the study and diagnosis of plant viral diseases in agriculture’. We deeply thank all colleagues from this COST action for their support to acquire the results - directly or indirectly - and for the fruitful and open scientific exchange.

## Supplementary Material

**Supplementary Table S1. Samples used for RT-PCR validation and genetic divergence analysis. Sample label, collection date, tree location, symptoms exhibited, damage caused by pests and the RT-PCR results (+/-) for each of the RT-PCR assays applied with seven different primer-pairs in 6 RNA segments of the novel emaravirus are shown**.

**Supplementary Table S2. Estimates of evolutionary divergence among RT-PCR products from 10 maple trees and the original Acer+ (2014). (A) RT-PCR products amplified with primer-pair RNA1aF/R. (B) RT-PCR products amplified with primer-pair RNAbF/R**.

